# Fbxw7 is a critical regulator of Schwann cell myelinating potential

**DOI:** 10.1101/342931

**Authors:** Breanne L. Harty, Fernanda Coelho, Sarah D. Ackerman, Amy L. Herbert, David A. Lyons, Kelly R. Monk

**Affiliations:** Department of Developmental Biology, Washington University School of Medicine; 660 S Euclid Ave, St Louis, MO 63110; United States of America; Vollum Institute, Oregon Health Sciences University; 3181 SW Sam Jackson Park Road, Portland, OR 97239; USA; Institute of Neuroscience, University of Oregon, Eugene, OR 97403; USA; Centre for Neuroregeneration, MS Society Centre for Translational Research, Euan MacDonald Centre for Motor Neurone Disease Research, University of Edinburgh, Edinburgh EH16 4SB, UK

**Keywords:** Schwann cell, Fbxw7, myelin, axon selection, myelinating potential

## Abstract

Myelin insulates and protects axons in vertebrate nervous systems. In the central nervous system (CNS), oligodendrocytes (OLs) make numerous myelin sheaths on multiple axons, whereas in the peripheral nervous system (PNS) myelinating Schwann cells (SCs) make just one myelin sheath on a single axon. Why the myelinating potentials of OLs and SCs are so fundamentally different is unclear. Here, we find that loss of Fbxw7, an E3 ubiquitin ligase component, enhances the myelinating potential of SCs. *Fbxw7* mutant SCs are seen myelinating multiple axons in a fashion reminiscent of OLs as well as aberrantly myelinating large axons while simultaneously ensheathing small unmyelinated axons - typically distinct roles of myelinating SCs and non-myelinating Remak SCs, respectively. We found that several of the *Fbxw7* mutant phenotypes, including the ability to generate thicker myelin sheaths, were due to dysregulation of mTOR. However, the remarkable ability of mutant SCs to either myelinate multiple axons or myelinate some axons while simultaneously encompassing other unmyelinated axons is independent of mTOR signaling. This indicates distinct roles for Fbxw7 in regulating multiple aspects of SC behavior and that novel Fbxw7-regulated mechanisms control modes of myelination previously thought to fundamentally distinguish myelinating SCs from non-myelinating SCs and OLs. Our data reveal unexpected plasticity in the myelinating potential of SCs, which may have important implications for our understanding of both PNS and CNS myelination and myelin repair.

## INTRODUCTION

In vertebrates, oligodendrocytes (OLs) and Schwann cells (SCs) are specialized glial cells that generate the myelin sheaths of the central nervous system (CNS) and peripheral nervous system (PNS), respectively. In both the CNS and PNS myelin enables fast action potential propagation and protects the axon that it surrounds [1]. Myelin sheaths of the CNS and PNS are broadly similar in composition and structure, and there is a large degree of overlap in the molecular control of myelination by OLs and myelinating SCs [2, 3]. However, there are important differences in how OLs and SCs interact with axons. One major fundamental difference between OLs and myelinating SCs is the ratio by which they myelinate axons. In peripheral nerves SCs select specific axons in a process called radial sorting, whereby they are thought to extend exploratory processes into a bundle of unmyelinated axons and select just one axon, greater than 1 µm in diameter, for myelination. Upon completion of radial sorting, promyelinating SCs are associated 1:1 with large-caliber axons (<1 µm in diameter), whereupon they initiate formation of a myelin sheath. In contrast, multiple small-caliber axons in peripheral nerves are ensheathed by non-myelinating Remak SCs into Remak bundles. The mechanisms controlling the Remak SC *vs.* myelinating SC fate are unclear. In the CNS, OLs also extend exploratory processes that dynamically interact with potential target axons [4], but unlike SCs, OLs are able to myelinate many axon segments [5]. Given the broad molecular similarities between myelinating SCs and OLs, it is unclear why SCs and OLs exhibit such differences in myelinating capacity.

The E3 ubiquitin ligase component *F-box and WD-repeat domain containing 7* (Fbxw7) is an important regulator of OL development and CNS myelination [6-8]. Here, using SC-specific knockout approaches in mice, we demonstrate that Fbxw7 plays distinct and novel roles in SCs. Of principal interest is that, in the absence of Fbxw7, SCs gain the ability to myelinate multiple axons in a fashion reminiscent of OLs. Also surprising was the ability of *Fbxw7* mutant SCs to generate myelin around large caliber axons while simultaneously ensheathing many additional small caliber axons. Electron microscopy and immunofluorescence (IF) analyses confirm that these cells are indeed SCs and not OLs that may have infiltrated the PNS. Additionally, *Fbxw7* mutants display early increases in SC number, smaller Remak bundles, and hypermyelination which are ameliorated upon loss of *mTOR*. However, even in the absence of mTOR, *Fbxw7* mutant SCs retain the remarkable ability to myelinate multiple axons, as well as simultaneously myelinate large axons while ensheathing small unmyelinated axons. This suggests that the molecular mechanisms that regulate the fundamental differences in myelinating potential between SCs and OLs are independent of mTOR signaling. Taken together, our findings show that the restriction of myelinating SCs to myelinate a single axon is not immutable and that Fbxw7 is a critical player in regulating the myelinating potential of SCs.

## RESULTS

### Fbxw7 cell-autonomously regulates SC development

Fbxw7 is a substrate recognition component of SKP1-Cullin-F-box (SCF) ubiquitin ligase complexes, which catalyze addition of ubiquitin moieties on certain proteins to target them for proteasomal degradation [9]. We and others have previously reported that *fbxw7* zebrafish mutants display striking overexpression of *myelin basic protein (mbp)* in the CNS, enhanced OL numbers and thicker CNS myelin [6-8].

A role for Fbxw7 in the PNS has never been described. However, given the role of Fbxw7 in OLs, we hypothesized that Fbxw7 is also required in SCs. To test this, we employed a conditional knockout strategy in mice. The *Dhh^cre^* transgene results in Cre recombinase expression under the *Desert hedgehog* promoter at approximately embryonic day (E) 12.5 in SC precursors [10]. To delete *Fbxw7* specifically in SCs, we crossed *Dhh^cre^* with an *Fbxw7^fl/fl^* transgenic line in which *loxP* sites flank exons 5 and 6 of *Fbxw7* [11]. This creates a frameshift upon Cre activity resulting in a null allele, which we confirmed by the absence of Fbxw7 mRNA by RT-PCR (Figure S1A). In all cases, *Dhh^cre(-)^;Fbxw7^fl/+^* and *Dhh^cre(-)^;Fbxw7^fl/fl^* siblings were used as controls.

We analyzed sciatic nerves by transmission electron microscopy (TEM) and found that at postnatal day 3 (P3), heterozygous *Dhh^cre(+)^;Fbxw7^fl**/+**^* (Het) and homozygous *Dhh^cre(+)^;Fbxw7^fl/fl^* (cKO) mutant mice displayed an increase in SC nuclei as well as in the proportion of myelinated axons relative to controls (Figure 1 A-E). However, by P42, SC numbers were equivalent in mutant and wild type nerves (Figure 1 F-J), suggesting that the early increase in SC number in *Fbxw7* mutants is transient. Nonetheless, myelin was considerably thicker in *Dhh^cre(+)^;Fbxw7^fl/+^* and *Dhh^cre(+)^;Fbxw^fl/fl^* mice compared to controls beginning at P21, especially on small caliber axons (Figure 1 K-N, Supplementary Figure 1 Q-R). Additionally, we found that Remak bundles contained fewer axons in *Dhh^cre(+)^;Fbxw7^fl/+^* and *Dhh^cre(+)^;Fbxw7^fl/fl^* nerves relative to controls (Figure S1 B-P). These data demonstrate that Fbxw7 functions cell-autonomously to regulate multiple aspects of SC development.

**Figure 1:**
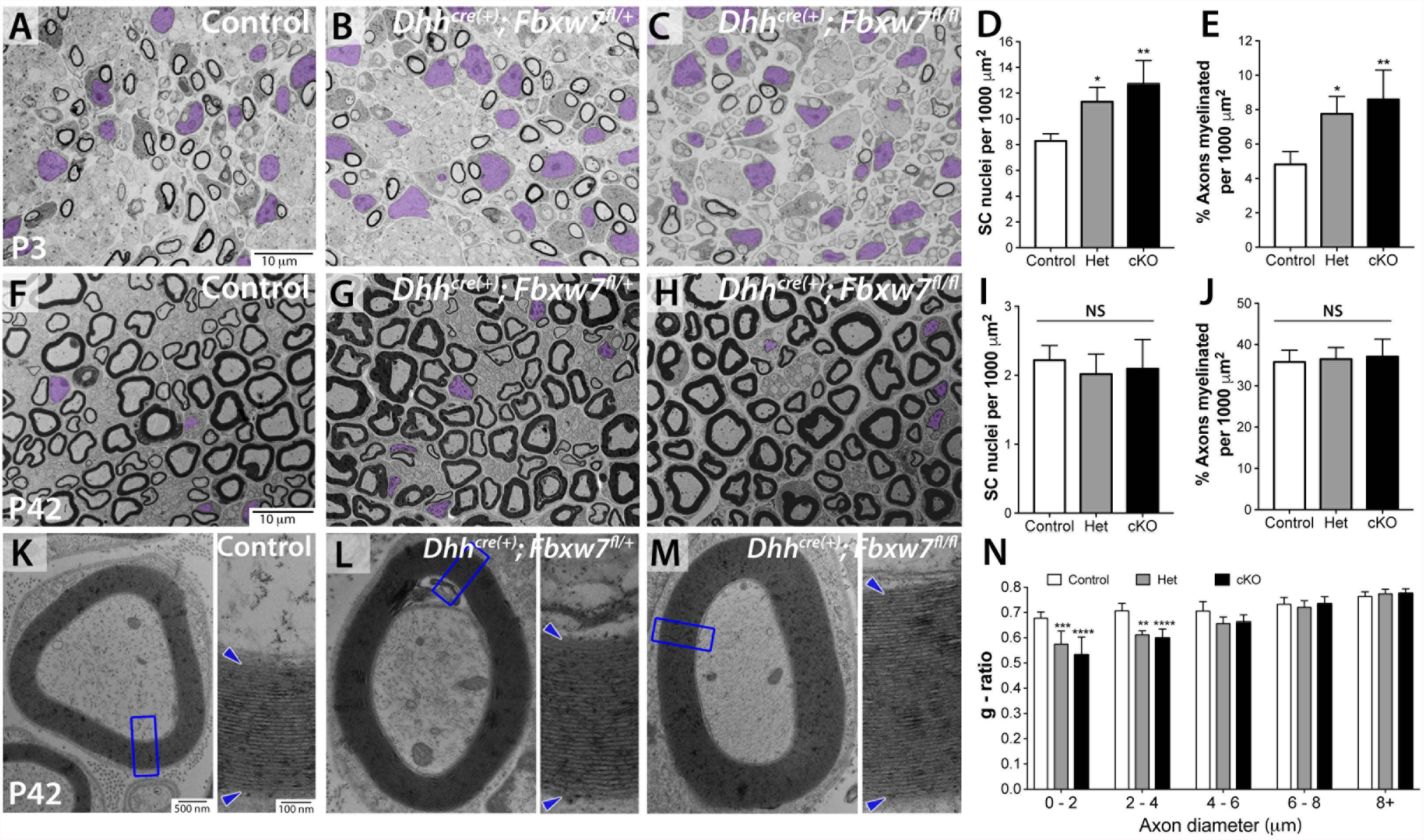
Fbxw7 is a conserved cell autonomous regulator of SC number and myelin thickness. (A-J) TEM of SC-specific *Fbxw7* mutant mice at P3 (A-E) and P42 (K-N). At P3, loss of *Fbxw7* leads to increased SC number (D; nuclei pseudocolored in purple) and percentage of myelinated axons (E) in both *Dhh^cre(+)^;Fbxw7^fl/+^* (Het) and *Dhh^cre(+)^;Fbxw7^fl/fl^* (cKO) nerves compared to littermate controls. However, by P42, both of these phenotypes have been resolved (I,J). Mutant SCs also make thicker myelin as evidenced by decreased g-ratios (K-M), especially on small diameter axons (N). Insets are zoomed images of the area indicated by blue boxes (U-W). Error bars depict S.D. ANOVA; * = p < 0.05, ** = p < 0.01, *** = p < 0.001. NS = not significant. Asterisks above bars indicate comparisons to controls. Comparisons between *Dhh^cre(+)^;Fbxw7^fl/+^* and *Dhh^cre(+)^;Fbxw7^fl/fl^* were not significant. P3: N = 4 control, 6 *Dhh^cre(+)^;Fbxw7^fl/+^*, 5 *Dhh^cre(+)^;Fbxw7^fl/fl^*. P42: N = 4 control, 3 *Dhh^cre(+)^; Fbxw7^fl/+^*, 4 *Dhh^cre(+)^; Fbxw7^fl/fl^*.

### Fbxw7 mutant Schwann cells can myelinate multiple axons

We also found that loss of Fbxw7 dramatically increased the myelinating potential of SCs (Figure 2, Supplementary Figure 2). Remarkably, in every *Dhh^cre(+)^;Fbxw7^fl/+^* and *Dhh^cre(+)^;Fbxw7^fl/fl^* nerve examined, even in adulthood, we observed numerous instances of multi-axonal myelination (Figure 2 B-C, F, I-J; Supplementary Figure 2 A-E). These myelinated axons were sometimes joined by long thin processes of SC cytoplasm in a manner reminiscent of OL myelination (Figure 2I, red arrow).

**Figure 2:**
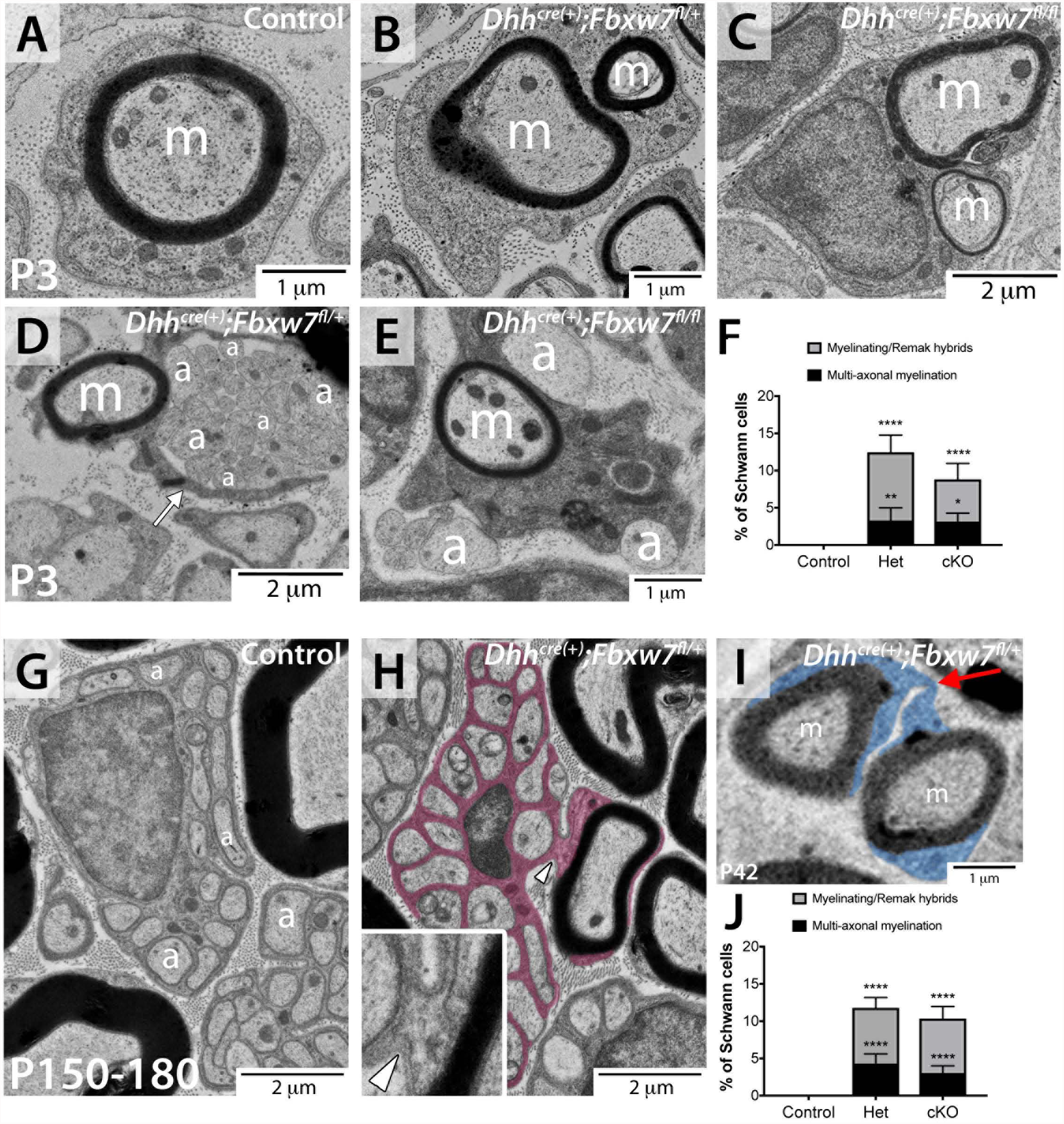
Loss of Fbxw7 enhances the myelinating capacity of SCs. TEM of SC-specific *Fbxw7* mutant mice show aberrant SC-axon interactions as early as P3 (A-F) and continuing until at least P180 (G-J). Loss of *Fbxw7* allowed SCs to myelinate multiple axons (B,C,I) as well as be able to both generate myelin as well as extend processes to encompass small caliber non-myelinated axons (D,E,H). These phenotypes were quantified together as aberrant SC-axon interactions, and were never observed in control nerves (F, J). In both cases there was continuous BL (white arrowhead) around these cells and their processes (H, inset), and it was the outer cytoplasmic pocket that extended these additional processes (D; white arrow). In aged nerves, some SCs still appeared to be myelinating/Remak “hybrids” (H; SC cytoplasm pseudocolored in red). The true proportion of mutant SCs displaying these behaviors is likely underestimated by cross-sectional analyses (F, J) as multiple myelinated axons were seen joined by very thin cytoplasmic processes (I; SC cytoplasm pseudocolored in blue). P3: N = 4 control, 6 *Dhh^cre(+)^;Fbxw7^fl/+^*, 5 *Dhh^cre(+)^;Fbxw7^fl/fl^*. P150-180: N = 5 control, 4 *Dhh^cre(+)^;Fbxw7^fl/+^*, 5 *Dhh^cre(+)^;Fbxw7^fl/fl^*. Error bars depict S.D. ANOVA; * = p < 0.05, ** = p < 0.01, *** = p < 0.001. Asterisks above bars indicate comparisons to controls; comparisons between *Dhh^cre(+)^;Fbxw7^fl/+^* and *Dhh^cre(+)^;Fbxw7^fl/fl^* were not significant. m = myelinated axon; a = unmyelinated axon.

We also observed *Fbxw7* mutant SCs that had myelinated one or more axons and simultaneously encompassed several unmyelinated axons in an immature SC or Remak SC-like fashion (Figure 2 D-F; Supplementary Figure 2 F-J). When mature, these SCs displayed both myelinating SC and Remak SC qualities almost as though these cells are myelinating/Remak SC “hybrids”, with the myelin being grossly normal and every unmyelinated axon being fully ensheathed by SC membrane and cytoplasm (Figure 2H, Supplementary Figure 2 H-J). Importantly, both of these distinct phenotypes were observed at all developmental stages examined – P3, P21, P42, and P150- 180 – suggesting that these are not simply transient developmental anomalies. It is also important to highlight that these phenotypes are distinct from “polyaxonal myelination” in which a bundle of axons is myelinated together as though it was one larger axon (Supplementary Figure 2K), which has been previously reported in mutants with radial sorting defects and occurs at low frequency even in wild-type nerves [12].

In total, by cross-sectional TEM analyses, approximately 12% of *Fbxw7* mutant SCs display enhanced myelinating potential, and this proportion remained consistent throughout the life of the animal (Figure 2 F, I). However, we found that in many cases multiple myelinated axons (Figure 2I; red arrow), or a myelinated axon and a bundle of unmyelinated axons (Supplementary Figure 2 G, J; blue arrow), are joined together by long thin processes of SC cytoplasm. This suggests that the proportion of SCs exhibiting enhanced myelinating potential may be underestimated by static cross-sectional imaging. To test this hypothesis, we performed serial block-face scanning electron microscopy to visualize portions of mutant nerves in 3D. We found that indeed, in most individual sections, myelinating cells appear to display the normal 1:1 SC:axon relationship. However, when we visualized cells in 3D it became clear that *Fbxw7* mutant SCs that look normal in one plane display phenotypes indicative of enhanced myelinating potential in other planes. For example, an *Fbxw7* mutant SC is observed myelinating multiple axons (magenta “m”) while simultaneously extending at least two long thin processes (yellow arrows and red box) to ensheath multiple unmyelinated axons (cyan “a”) (Supplemental Movie 1).

Importantly, in both types of aberrant SC-axon interactions in *Fbxw7* mutants, we were able to trace continuous SC cytoplasm and basal lamina on the abaxonal surface of the mutant SC (Figure 2H, inset, white arrowhead). Since SCs secrete a basal lamina and OLs do not [13], the presence of a basal lamina strongly supports the notion that these cells are indeed SCs and not OLs that might have infiltrated the PNS. To further confirm that the mutant cells are not OLs, we analyzed *Fbxw7* mutant sciatic nerves by immunofluorescence for markers of the OL lineage. We did not find any evidence of Olig2-positive cells (immature OL lineage cells) in either control or *Fbxw7* mutant nerves (data not shown). These data suggest that the enhanced myelinating potential observed in *Fbxw7* mutants is not due to OL infiltration. Instead, it suggests that mutant SCs are extending multiple cytoplasmic projections and that more than one of these processes can go on to make myelin.

### Fbxw7 regulates mTOR to control SC number, myelination, and Remak bundle organization

To our knowledge, no genetic or pharmacological manipulation *in vivo* has been reported to increase the myelinating potential of SCs in the manner observed in *Fbxw7* mutants. However, the hypermyelination and oversorted Remak bundle defects observed in *Fbxw7* mutants have also been described following SC-specific deletion of *Pten* [14] or overactivation of *Akt* [15], which both result in elevated mTOR signaling. Previous reports show loss of Fbxw7 function enhances levels of mTOR and its targets [16], and Fbxw7 was recently shown to regulate mTOR in CNS myelination [7].

Therefore, we examined levels of mTOR and found that total mTOR protein levels are approximately 2-fold higher in *Fbxw7* mutants relative to controls (Figure 3A-B). Similarly, consistent with previous findings that multiple feedback loops regulate the PI3K/mTOR pathway [17], we found that the mRNA levels of mTOR as well as several targets of mTOR were also significantly elevated in *Fbxw7* mutant nerves (Figure 3C).

**Figure 3:**
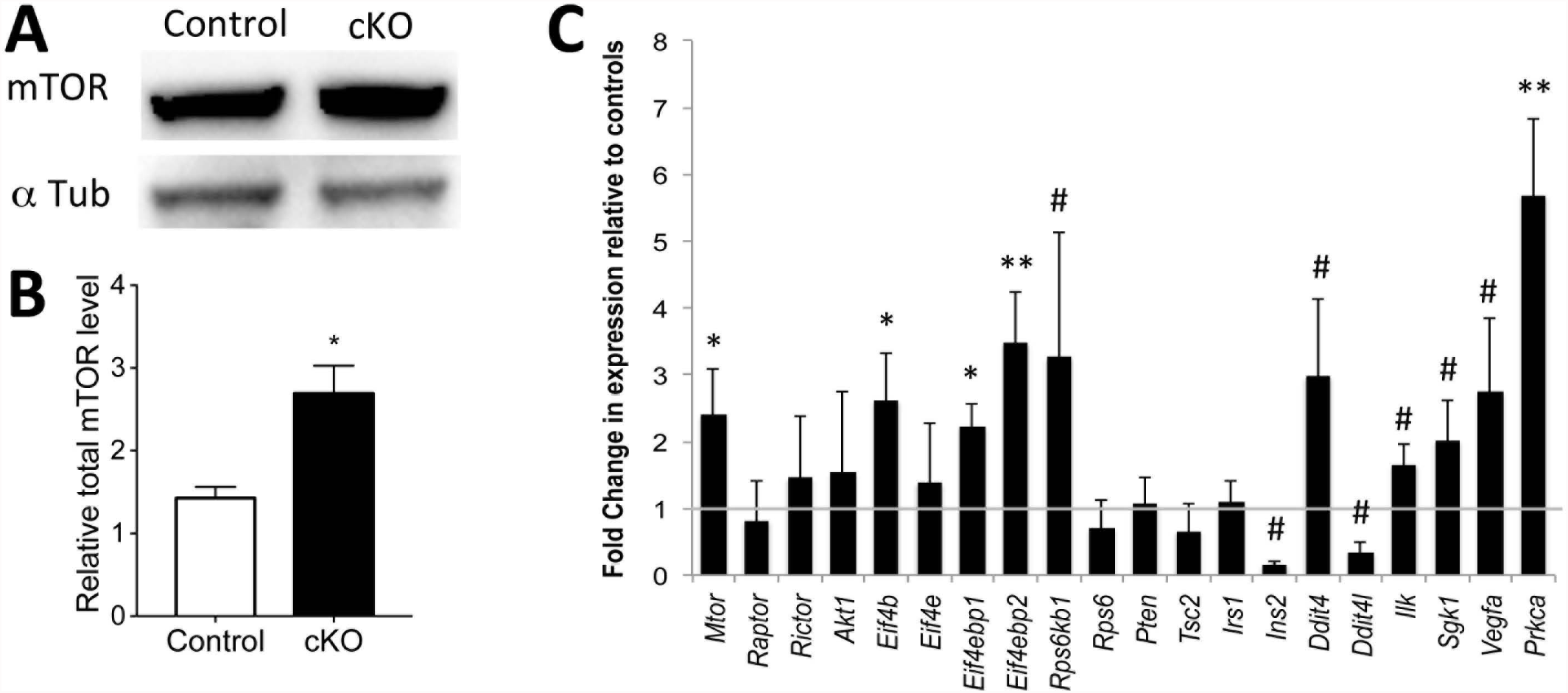
mTOR levels are elevated in *Fbxw7* mutants. Western blot analyses of sciatic nerve lysates indicate a 2-fold increase in mTOR protein in *Dhh^cre(+)^:Fbxw7^fl/fl^* (cKO) nerves compared to littermate controls when normalized to both background and alpha-tubulin levels (A, B). *mTOR* mRNA is also upregulated in *Dhh^cre(+)^:Fbxw7^fl/fl^* nerves (C), as are multiple mTOR targets. The gray line at y = 1 denotes control levels. For western blots: N = 3 controls, N = 3 *Dhh^cre(+)^:Fbxw7^fl/fl^* at P42. For qPCR, N = 3 control and 3 *Dhh^cre(+)^:Fbxw7^fl/fl^* at P21. Error bars depict S.D. ANOVA; # = p < 0.1, * = p < 0.05,**=p < 0.01.

To directly test if Fbxw7 controls mTOR in SCs, we generated double SC-specific *Fbxw7;mTOR* knockouts by crossing *mTOR^fl/fl^* mice [18] to our *Dhh^cre(+)^;Fbxw7^fl/fl^* mice (Supplementary Figure 3 and Figure 4). We were unable to obtain enough *Dhh^cre(+)^;Fbxw7^fl/fl^;mTOR^fl/fl^* mutants from trans-heterozygous intercrosses (1/32 expected, 1/126 obtained (Figure 4 E,E’)); therefore, we analyzed *Dhh^cre(+)^;Fbxw7^fl/+^;mTOR^fl/fl^* (HetΔmTOR) animals in more detail as the *Fbxw7* mutation is dominant. For the epistasis experiments, *Dhh^cre(-)^;Fbxw7^fl/+^;mTOR^fl/+^* and *Dhh^cre(-)^;Fbxw7^fl/+^;mTOR^fl/fl^* siblings were used as controls.

**Figure 4:**
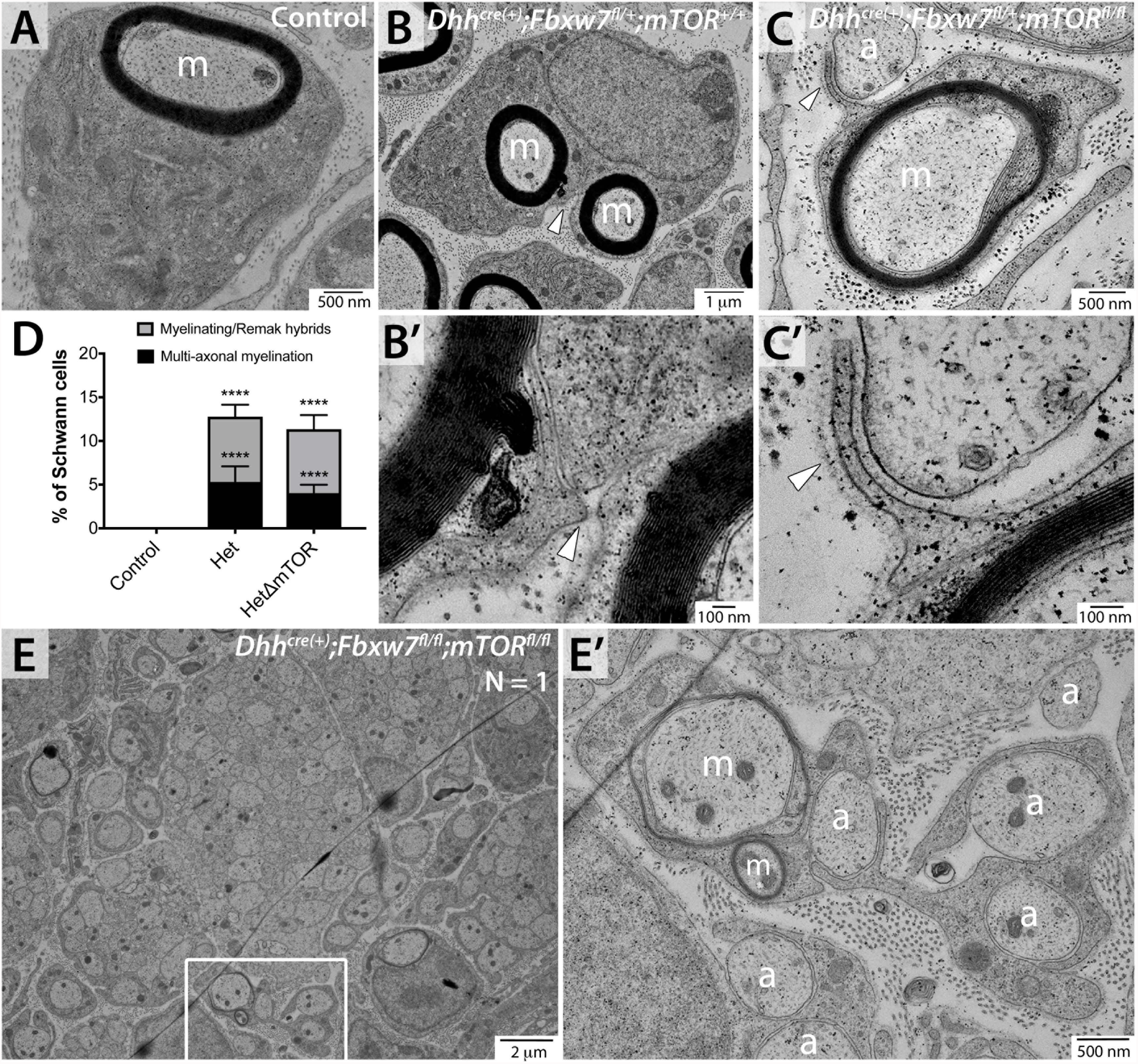
The myelinating capacity of SCs is independent of mTOR signaling. *mTOR* deletion in *Fbxw7* mutant nerves failed to suppress the aberrant SC-axon interactions. Multi-axonal myelination and SCs that had both myelinated and encompassed multiple small non-myelinated axons were present in both *Dhh^cre(+)^;Fbxw7^fl/+^;mTOR^+/+^* (B, B’) and *Dhh^cre(+)^;Fbxw7^fl/+^;mTOR^fl/fl^* (HetΔmTOR) (C, C’) nerves. Neither phenotype was ever observed in sibling controls (D). Tracing the BL (B’, C’; white arrows) confirmed these behaviors were the acts of single SCs. Both phenotypes were also present in the sole *Dhh^cre(+)^;Fbxw7^fl/fl^;mTOR^fl/fl^* (cKOΔmTOR) animal we were able to collect (E, E’). Error bars depict SD. ANOVA; *** = p < 0.001. Asterisks above bars indicate comparisons to controls; comparisons between *Dhh^cre(+)^;Fbxw7^fl/+^* and *Dhh^cre(+)^;Fbxw7^fl/+^;mTOR^fl/fl^* were not significant. P3: N = 4 control, 6 *Dhh^cre(+)^;Fbxw7^fl//+^;mTOR^+/+^*, 5 *Dhh^cre(+)^;Fbxw7^fl/+^;mTOR^fl/fl^*, 1 *Dhh^cre(+)^;Fbxw7^fl/fl^;mTOR^fl/fl^*. m = myelinated axon; a = unmyelinated axon.

If mTOR is the Fbxw7 target responsible for the SC defects observed in *Fbxw7* mutants, loss of *mTOR* should suppress these phenotypes in *Dhh^cre(+)^;Fbxw7^fl/+^* animals. Indeed, at P3, *Dhh^cre(+)^;Fbxw7^fl/+^;mTOR^fl/fl^* animals were indistinguishable from controls in SC numbers and the percentage of myelinated axons (Supplementary Figure 3 A-E). At P21, Remak bundles in *Dhh^cre(+)^;Fbxw7^fl/+^;mTOR^fl/fl^* nerves contained significantly more axons relative to Remak bundles in either controls or *Fbxw7* mutants (Supplementary Figure 3 F-J). Further, in agreement with previous studies [19], *Dhh^cre(+)^;Fbxw7^fl/+^;mTOR^fl/fl^* nerves were hypomyelinated relative to controls and *Dhh^cre(+)^;Fbxw7^fl/+^* animals (Supplementary Figure 4K), demonstrating that *mTOR* is epistatic to *Fbxw7*. These data suggest that Fbxw7 regulates mTOR to control early SC number, myelin thickness, and axon ensheathment by Remak SCs.

### SC myelinating potential is independent of mTOR signaling

Strikingly, however, loss of *mTOR* in *Fbxw7* mutant SCs was unable to restore normal SC myelinating potential. Despite the fact that *Dhh^cre(+)^;Fbxw7^fl/+^;mTOR^fl/fl^* nerves had normal numbers of SCs, delayed radial sorting, larger Remak bundles, and thinner myelin, loss of mTOR was insufficient to suppress the enhanced myelinating potential observed in *Fbxw7* mutant SCs. Double mutant SCs were still seen myelinating multiple axons, as well as displaying myelinating/Remak “hybrid” phenotypes (Figure 4). As in the *Dhh^cre(+)^;Fbxw7^fl/+^* and *Dhh^cre(+)^;Fbxw7^fl/fl^* animals, mutant SCs in *Dhh^cre(+)^;Fbxw7^fl/+^;mTOR^fl/fl^* and *Dhh^cre(+)^;Fbxw7^fl/fl^;mTOR^fl/fl^* nerves that had already myelinated large axons were seen extending processes from the outer cytoplasmic pocket to interact with other axons (Figure 4 C, C’; white arrowhead). These data suggest that Fbxw7 plays a novel role in controlling SC myelinating potential that is independent of mTOR signaling (Figure 5). It is remarkable that SC myelinating capacity remains enhanced in *Dhh^cre(+)^;Fbxw7^fl/+^;mTOR^fl/fl^* mutants despite the fact that deletion of *mTOR* has caused reduced myelin thickness, delayed radial sorting, and defects in Remak SC ensheathment. This suggests that the morphological processes controlling the myelinating potential of SCs are distinct from the cellular behaviors involved in radial sorting, myelination, or Remak SC ensheathment.

**Figure 5:**
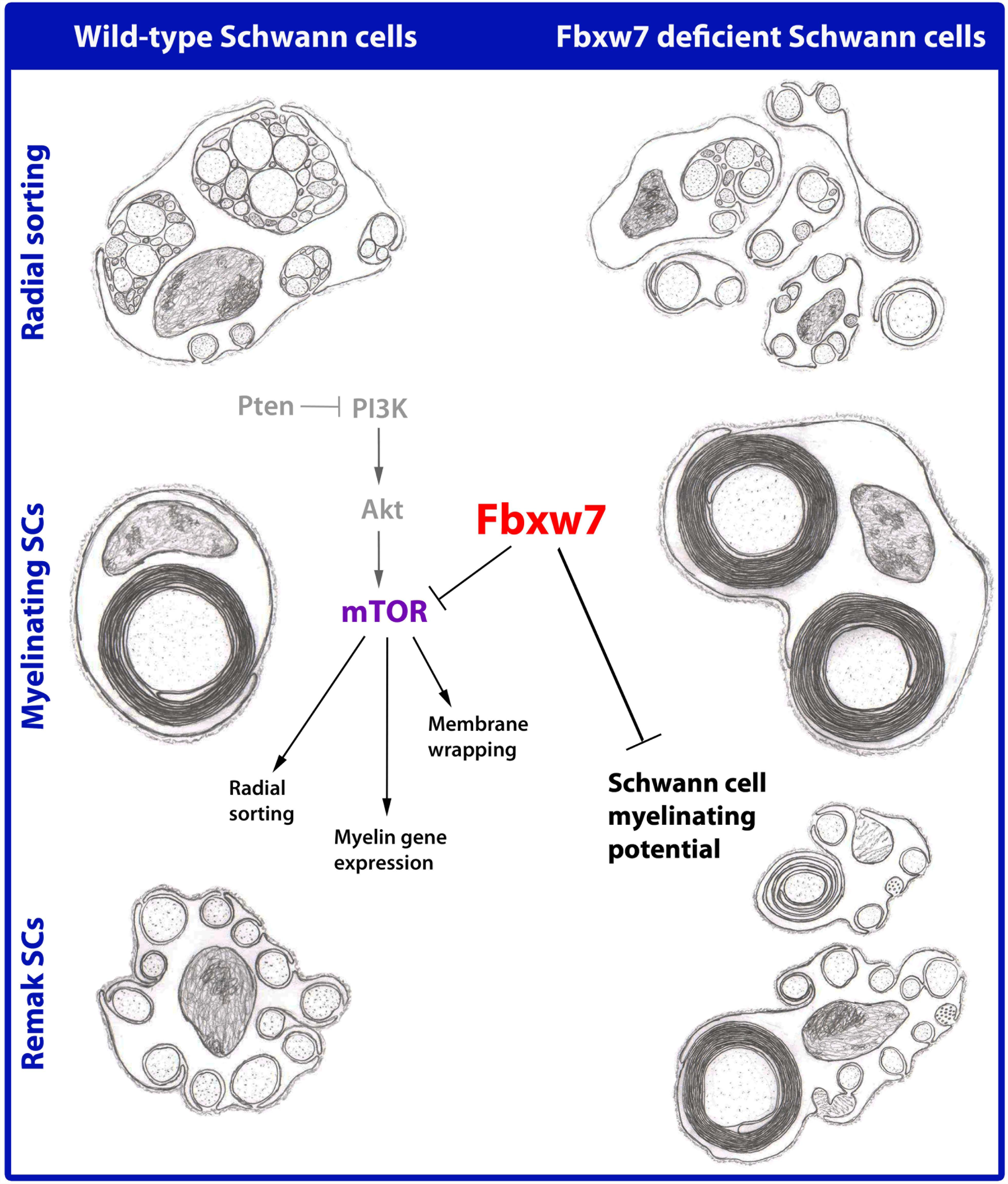
Fbxw7 orchestrates SC biology via mTOR-dependent and –independent mechanisms. Artistic renditions of the phenotypes observed in *Fbxw7* mutant SCs (right) as compared to normal SCs (left). Fbxw7 is involved in radial sorting (top), as well as both mature myelinating SCs (middle) and Remak SCs (bottom). A simplified PI3K/mTOR pathway shows that in SCs, Fbxw7 directly inhibits mTOR, and thus regulates multiple aspects of SC biology. Fbxw7 also targets many other proteins that are essential for various cellular processes, and one of these may be responsible for controlling the myelinating capacity of SCs. Alternatively, it is possible that this is a novel role of Fbxw7 independent of its function in E3 ubiquitin ligase complexes. Steps in the pathway that were not demonstrated directly in this study are shown in gray. Pencil sketches by BLH.

## DISCUSSION

In stark contrast to OLs in the CNS, which can myelinate dozens of axons simultaneously, myelinating SCs are restricted to myelinating a single axonal segment in the PNS. The molecular mechanisms controlling the differences in myelinating potential between SCs and OLs remain mysterious. Here we show that SCs are capable of myelinating multiple axons *in vivo* and challenge the notion that the ability of a SC to extend multiple processes is mutually exclusive with the capacity to make myelin. Remak SCs are also capable of extending multiple processes and interacting with many axons, but unlike OLs they do not myelinate axons. However, upon loss of *Fbxw7*, single SCs gain the ability to both myelinate some axons, as well as encompass many unmyelinated axons as if *Fbxw7* mutant SCs are myelinating SC and Remak SC “hybrids.” The multi-axonal myelination observed in *Fbxw7* mutant SCs is reminiscent of an OL’s ability to simultaneously extend multiple cytoplasmic processes and myelinate multiple axon segments. However, the presence of basal lamina and lack of OL-lineage markers suggests that OLs have not infiltrated *Fbxw7* mutant nerves. Perhaps Fbxw7 plays a role in inhibiting the myelinating potential of SCs by limiting the extent to which they can generate multiple processes, thus restricting myelinating SCs to a single axonal segment.

Although neither of the aberrant *Fbxw7* mutant SC-axon interaction phenotypes have been previously described *in vivo*, several of the other phenotypes observed in *Fbxw7* mutant nerves resemble phenotypes described in mutants where mTOR signaling is enhanced such as in *Pten* mutants [14] and constitutively active Akt mutants [15]. It is well documented, from these studies and others, that mTOR levels must be tightly regulated in SCs such that any type of manipulation results in defective PNS myelination [14, 15, 19-25]. mTOR is a bona fide target of Fbxw7 in other contexts [16] and Fbxw7 was recently shown to control OL myelination through mTOR [7]. Thus, we tested the hypothesis that Fbxw7 regulates mTOR to control SC development. We showed that mTOR and some of its targets are upregulated in *Fbxw7* mutants, suggesting that mTOR activity is elevated. Double transgenic analysis demonstrated that *mTOR* is epistatic to *Fbxw7* and is responsible for regulating early SC numbers, appropriate myelin thickness, and Remak bundle organization.

Notably, however, loss of *mTOR* in *Fbxw7* mutant SCs was unable to restore typical SC:axon ratios. Despite the fact that *Dhh^cre(+)^;Fbxw7^fl/+^;mTOR^fl/fl^* nerves had normal numbers of SCs, delayed radial sorting, larger Remak bundles, and thinner myelin, loss of *Fbxw7* was nevertheless sufficient to drive increased myelinating potential of SCs. This suggests that the myelinating potential of SCs is independent of mTOR signaling, and importantly, that the mechanisms controlling the typical SC:axon ratio is also controlled independently from other morphological behaviors of SCs, including radial sorting, Remak SC ensheathment, and membrane wrapping.

This highlights the complicated nature of removing a gene like Fbxw7, which is a master regulator of master regulators. Given the role of Fbxw7 in E3 ubiquitin ligase complexes, it is likely that there is an as yet unidentified target protein that is dysregulated in *Fbxw7* mutants that is responsible for the enhanced myelinating potential. However, it is equally possible that the complex phenotypes observed in Fbxw7 mutant nerves result from combinatorial interactions amongst multiple misregulated targets, or that this represents a novel function of Fbxw7.

A potentially interesting future direction would be to explore the hypothesis that *Fbxw7* mutant SCs are behaving similarly to repair SCs. After PNS injury, SCs readily become repair SCs, which clear axon and myelin debris, recruit macrophages, aid in axon regrowth, and finally remyelinate axons [26]. A recent study on the length and morphology of SCs throughout development, differentiation, and remyelination found that repair SCs become extremely long and branched after injury [27]. Given the observation that myelinated axons were joined by long thin processes of SC cytoplasm in *Fbxw7* mutants it is possible that *Fbxw7* mutant SCs have also assumed a branched morphology. Although dispensable during SC development, the key switch controlling the transition to the repair SC phenotype is upregulation of the transcription factor c-Jun [28, 29]. Importantly, c-Jun is a well-known target of Fbxw7 in other contexts [30], and loss of Fbxw7 has previously been shown to elevate c-Jun levels [31]. Recent evidence also suggests that mTOR is transiently reactivated after nerve injury to promote the elevation of c-Jun [21]. Thus, future efforts should assess c-Jun levels in *Fbxw7* mutant nerves and explore the implications of enhanced myelinating potential on remyelination after PNS injury.

In contrast to SC remyelination, OL remyelination and CNS recovery after injury is limited in mammals, and OLs do not de/redifferentiate to aid in recovery [32]. One interesting hypothesis is that the distinction between the ability of SCs *vs*. OLs to facilitate repair after injury is rooted in fundamental qualities that distinguish these cells, such as the number of axons they associate with and myelinate. It is now clear that OLs interpret and respond to neuronal activity by selective myelination and that OLs play a critical role in circuit development and/or maintenance [33-36]. It is intriguing to speculate that OLs may link circuits through their interactions with and myelination of multiple axons. However, in the PNS, where circuits are less complex, there is not strong evidence for SC participation in circuit function. Rather, more emphasis seems to be on the ability of SCs to rapidly and faithfully respond to injuries, which occur more readily in the PNS. Presumably this rapid response would be more difficult if SCs were required to demyelinate more than one axon, especially if only some of those axons were injured, while others were intact. Given that *Fbxw7* mutant SCs provide the first *in vivo* description of multi-axonal myelination by SCs, *Fbxw7* mutants represent a unique and useful tool with which to investigate the impact of differences between myelinating SCs, Remak SCs, and OLs on nervous system repair.

## AUTHOR CONTRIBUTIONS

Conceptualization and Methodology, all authors; Investigation, BLH and FC; Formal Analysis, BLH and FC; Writing – Original Draft, BLH, DAL, and KRM; Writing – Review and Editing, all authors; Visualization, BLH and FC; Funding Acquisition, KRM.

## ACKNOWLEDGEMENTS

The authors thank the Monk lab (Washington University [WU] and Oregon Health & Science University [OHSU]) for valuable discussions, as well as the Solnica-Krezel and Johnson laboratories at Washington University (WU) for helpful feedback. The Cavalli Lab (WU), especially Dan Carlin, supplied helpful reagents and useful feedback. We also thank the Nechiporuk Lab at OHSU for helpful input and advice. We are indebted to Laura Feltri, Carmen Melendez-Vasquez, and Steve Scherer for helpful discussions and to Megan Corty and Ben Emery for critical comments on the manuscript. This work was supported by: NIH/NINDS to BLH (F31 NS094004); NIH/NINDS to SDA (F31 NS087801); NIH/NINDS to ALH (F31 NS096814); the Edward J. Mallinckrodt Jr. Foundation (KRM), a Wellcome Senior Research Fellowship (DAL), and KRM is a Harry Weaver Neuroscience Scholar of the National Multiple Sclerosis Society.

## FIGURE LEGENDS

**Supplementary Figure 1: Fbxw7 cell-autonomously regulates proper SC radial sorting and Remak bundle ensheathment.** Schematic of deletion of exons 5 and 6 of Fbxw7 in *Dhh^cre(+)^:Fbxw7^fl/fl^* animals (A). RT-PCR shows that *Fbxw7* mRNA is significantly reduced in sciatic nerves of *Dhh^cre(+)^/Fbxw7^fl/fl^* (cKO) compared to *Dhh^cre(-)^/Fbxw7^fl/fl^* (control) at P21 (A). Ultrastructural analyses at P3 show that loss of *Fbxw7* results in increased segregation of axons during radial sorting (P3; B-F), as measured by decreased average number of axons/bundle (E), as well as the percentage of bundles having 5 or fewer axons (F). This phenotype persists at P21 (G-K) and until at least 6 months of age (P150-P180; L-P). Myelin thickness is increased, particularly on small axons, in *Fbxw7* mutant nerves beginning at P21 (Q) and persisting until at least P180 (R). Error bars depict S.D. ANOVA; * = p < 0.05, ** = p < 0.01, *** = p < 0.001, **** = p < 0.0001, NS = not significant. Asterisks above bars indicate comparisons to controls. Unless otherwise indicated, comparisons between *Dhh^cre(+)^;Fbxw7^fl/+^* and *Dhh^cre(+)^;Fbxw7^fl/fl^* were not significant. P3: N = 4 control, 6 *Dhh^cre(+)^:Fbxw7^fl/+^* (Het), 5 *Dhh^cre(+)^:Fbxw7^fl/fl^*. P21: N = 3 control, 4 *Dhh^cre(+)^:Fbxw7^fl/+^*, 4 *Dhh^cre(+)^: Fbxw7^fl/fl^*. P150-180: N = 5 control, 4 *Dhh^cre(+)^: Fbxw7^fl/+^*, 5 *Dhh^cre(+)^: Fbxw7^fl/fl^*.

**Supplementary Figure 2: Enhanced myelinating potential in *Fbxw7* mutant SCs.** Additional examples of multi-axonal myelination in Het and cKO animals at each time point – P3 (A,F), P21 (BD, G), P42 (E, H, I) and P150-180 (J). *Fbxw7* mutant SCs were occasionally seen myelinating as many as four independent axons (A; myelinated axons indicated by “m”; unmyelinated axons indicated by “a”). (A, F-J) Additional examples of SCs that have both myelinated larger axons as well as encompassed several non-myelinated axons like myelinating/Remak SC “hybrids” – P3 (A, F), P21 (G), P42 (E, H, I) and P150-180 (J). Here again, the myelinated axon and the bundle of unmyelinated axons were sometimes linked by thin processes of SC cytoplasm (G, J; blue arrows). These phenotypes are distinct from “polyaxonal myelination” in which a bundle of small-diameter axons is myelinated as though the bundle were a single large axon (K).

**Supplementary Figure 3: Genetic inhibition of mTOR suppresses the SC number, radial sorting, and myelin thickness phenotypes seen in Fbxw7 mutant nerves.** (A-J) TEM of SC-specific double mutant *Fbxw7:mTOR* mice at P3 (A-E) and P21 (F-J) demonstrates that *Dhh^cre(+)^;Fbxw7^fl//+^;mTOR^+/+^* (B; Het) nerves display greater numbers of SC nuclei as well as a higher percentage of myelinated axons relative to controls (A). Loss of mTOR in *Dhh^cre(+)^;Fbxw7^fl/+^;mTOR^fl/fl^* (HetΔmTOR; C) nerves suppresses both of these phenotypes (D, E). Deletion of *mTOR* further suppressed the radial sorting and Remak SC ensheathment defects observed in *Dhh^cre(+)^;Fbxw7^fl//+^;mTOR^+/+^* nerves (F-J). *Dhh^cre(+)^;Fbxw7^fl/+^;mTOR^fl/fl^* sciatic nerves also displayed markedly thinner myelin (increased g-ratios) relative to both control and *Dhh^cre(+)^;Fbxw7^fl//+^;mTOR^+/+^* siblings (K). P3: N = 4 controls, N = 6 *Dhh^cre(+)^;Fbxw7^fl//+^;mTOR^+/+^*, N = 5 *Dhh^cre(+)^;Fbxw7^fl/+^;mTOR^fl/fl^*. For P21: N = 6 controls, N = 4 *Dhh^cre(+)^;Fbxw7^fl//+^;mTOR^+/+^*, and N = 3 *Dhh^cre(+)^;Fbxw7^fl/+^;mTOR^fl/fl^*. Error bars depict S.D. One-way ANOVA for all measures except g-ratio, where we used two-way ANOVA in order to compare by both genotype and axon diameter; * = p < 0.05, ** = p < 0.01, *** = p < 0.001, **** = p < 0.0001.

**Supplemental Movie 1: 3D Imaging of *Fbxw7* mutant nerves.** Serial block-face scanning electron microscopy was used to image an *Fbxw7* mutant nerve at P180. Despite the fact that most individual sections show mutant SCs appearing with normal SC:axon ratios, when sections are imaged in sequence, it is clear that *Fbxw7* mutant SCs display enhanced myelinating potential. In the beginning of the movie, the cell pictured is seen myelinating two different axons (magenta “m”). Later, the SC cytoplasm around a bundle of unmyelinated axons (cyan “a”) in the lower left corner of the field is seen extending two long thin cytoplasmic processes (yellow arrows). One of these processes eventually joins the unmyelinated bundle with the myelinating SC (red box), indicating that the same SC is both myelinating some axons while ensheathing other unmyelinated axons at the same time.

## MATERIALS AND METHODS

### CONTACT FOR REAGENT AND RESOURCE SHARING

Further information and requests for resources and reagents should be directed to and will be fulfilled by Kelly Monk.

### EXPERIMENTAL MODEL AND SUBJECT DETAILS

All animal experiments were performed in compliance with Washington University’s institutional animal protocols.

The Fbxw7 conditional-ready mice (*Fbxw7^fl/fl^*) [11] were obtained from Jackson laboratories (Stock #: 017563) on a pure C57BL/6 background. *Fbxw7^fl/fl^* mice were mated to *Dhh^Cre(+)^* mice [10] that had also been maintained as pure on C57BL/6 (> 7 generations) to generate *Dhh^Cre(+)^;Fbxw7^fl/+^* (Het) mice. *Fbxw7* Hets were backcrossed to *Fbxw7^fl/fl^* animals to obtain *Dhh^Cre(+)^;Fbxw7^fl/f l^*(cKO) mice. For all cKO experiments, we used *Dhh^Cre(-)^;Fbxw7^fl/+^* or *Dhh^Cre(-)^;Fbxw7^fl/fl^* littermates as controls. For the double mutant experiments with Fbxw7 and mTOR, we obtained mTOR conditional-ready (*mTOR^fl/fl^*) mice [37] from Jackson laboratories (Stock #: 0110009), also on a pure on C57BL/6 background. The *mTOR^fl/fl^* mice were crossed with *Dhh^Cre(+)^;Fbxw7^fl/fl^* animals to generate *Dhh^Cre(+)^;Fbxw7^fl/+^;mTOR^fl/+^* animals. Finally, to obtain double mutants, we crossed *Dhh^Cre(+)^;Fbxw7^fl/+^;mTOR^fl/+^* to *Dhh^Cre(-)^;Fbxw7^fl/+^;mTOR^fl/+^* animals and analyzed *Dhh^Cre(+)^;Fbxw7^fl/+^;mTOR^fl/fl^* animals (“HetΔmTOR” for brevity). In all cases, mice of both sexes were analyzed, in equal ratios whenever possible. In all cases mutants were compared with littermate sibling controls.

All mouse lines were genotyped as previously described [10, 11, 37].

## METHOD DETAILS

### Transmission electron microscopy (TEM)

TEM was performed on mouse sciatic nerves at P3, P21, P42 and 6 months as previously described [38]. Briefly, nerves were drop-fixed in modified Karnovsky’s fixative (4% PFA, 2% glutaraldehyde, 0.1M sodium cacodylate, pH 7.4) at least overnight at 4**°**C. Samples were then washed with 0.1M sodium cacodylate to remove fixative, and then post-fixed for 1 hour in 2% osmium tetroxide in 0.1M sodium cacodylate. Nerves were then dehydrated with increasing concentrations of ethanol followed by propylene oxide (PO). Samples were then infiltrated for 1-2 hours in 2:1 PO:EPON, and then overnight in 1:1 PO:EPON with gentle agitation at room temperature. Samples were then transferred to 100% EPON while residual PO was allowed to fully evaporate (>4 hours). Sectioning, grid staining, imaging, and image analyses were performed as described for zebrafish. For all time points, four non-overlapping images at 1000X magnification were quantified, and all mouse TEM data is expressed per 1000 µm^2^ area. Control genotypes were: *Dhh^Cre(^*^-)^*;Fbxw7^fl/+^* and *Dhh^Cre(-)^;Fbxw7^fl/fl^*. For P3: N = 4 controls, N = 6 *Dhh^Cre(^*^+)^*;Fbxw7^fl/+^* (Hets), and N = 5 *Dhh^Cre(^*^+)^*;Fbxw7^fl/fl^* (cKO). At P21: N = 4 controls, N = 4 Hets, and N = 4 cKO. For P42 samples: N = 4 controls, N = 3 Hets, and N = 4 cKO. At 6 months of age: N = 5 controls, N = 4 Hets, N = 5 cKO. For the double mutant analyses, control genotypes used were: *Dhh^Cre(^*^-)^*;Fbxw7^fl/+^;mTOR^+/+^ Dhh^Cre(^*^-)^*;Fbxw7^fl/+^;mTOR^fl/fl^, Dhh^Cre(^*^-)^*;Fbxw7^fl/+^;mTOR^fl/+^*, *Dhh^Cre(^*^-)^*;Fbxw7^fl/fl^;mTOR^fl/fl^*, and *Dhh^Cre(^*^-)^*;Fbxw7^+/+^;mTOR^fl/fl^*. Fbxw7 “Hets” were *Dhh^Cre(^*^+)^*;Fbxw7^fl/+^;mTOR^+/+^* littermate siblings, and “HetΔmTOR” animals were *Dhh^Cre(^*^+)^*;Fbxw7^fl/+^;mTOR^fl/fl^* siblings. At P3: N = 4 controls, N = 6 Hets, N = 5 HetΔmTOR, N = 1 *Dhh^Cre(^*^+)^*;Fbxw7^fl/fl^;mTOR^fl/fl^* (cKOΔmTOR). For P21: N = 6 controls, N = 4 Hets, and N = 3 HetΔmTOR.

### Serial block-face scanning electron microscopy (SBF-SEM)

*<u>SBF-SEM was performed on a Dhh^Cre(^</u>*<u>^+)^*;Fbxw7^f^*</u>*^l/fl^* nerve at P180. Nerves were fixed in 4% PFA overnight at 4**°**C. Nerves were then processed for SBF-SEM by the Multiscale Microscopy Core at Oregon Health & Science University. Images were acquired every 50 nm using an FEI Teneo VolumeScope Microscope. Sections were annotated using FIJI and then the movie was composed using Microscopy Image Browser software.

### Western blot analyses

To assess mTOR protein levels in the sciatic nerve, we dissected nerves from the sciatic notch to just proximal to the trifurcation. These nerve segments were flash frozen in liquid nitrogen, cut into small pieces with microdissection scissors, and homogenized in lysis buffer (20 mm Tris-HCl, pH 7.5, 150 mm NaCl, 1 mm Na2EDTA, 1 mm EGTA, 1% Triton X-100, 2.5 mm sodium pyrophosphate, 1 mm β-glycerophosphate, 1 mm Na3VO4, 1 μg/ml leupeptin) with phosphatase inhibitor mixtures 1 and 2 (Invitrogen, ThermoFisher Scientific, Waltham, MA). Equal protein amounts (15 μg) were loaded and analyzed by SDS-PAGE and western blot. Antibodies used were: anti-mTOR (1:1000; Cell Signaling Technologies, Danvers, MA), and anti-α-tubulin (1:1000; Abcam, Cambridge, MA). Western blot images were quantified using FIJI, all bands were normalized to background and mTOR bands were compared to α-tubulin levels.

### RNA isolation and reverse transcription

Total RNA was extracted from flash-frozen P21 mouse sciatic nerves (N=3 *Dhh^Cre(^*^-)^*;Fbxw7^fl/fl^* littermate controls and N=3 *Dhh^Cre(^*^+)^*;Fbxw7^fl/fl^* animals), using a standard TRIzol extraction protocol (Life Technologies, ThermoFisher Scientific, Waltham, MA). Briefly, TRIzol was added to the frozen tissue samples, which were then allowed to thaw at room temperature for 10 min. During this incubation time, and while still in TRIzol, nerves were cut into much smaller pieces using microdissection scissors. Samples were homogenized via disruption with a plastic-tipped electric homogenizer, followed by passage through syringe and 22.5g needle at least ten times, and then a 27g needle at least ten more times until no lumps of tissue were observed. Once the nerves had been homogenized, we proceeded as usual with the standard TRIzol RNA extraction procedure as per manufacturer instructions.

Total RNA (500 ng) was then reverse transcribed in 20 μl using the Superscript III First Strand cDNA Synthesis Kit (Invitrogen, ThermoFisher Scientific, Waltham, MA) using random hexamers, as per manufacturer instructions. All cDNA products were diluted 1:5 prior to use in qPCR reactions.

### Quantitative reverse transcription PCR

To assay mRNA expression levels of mTOR and members of the mTOR signaling pathway, we used the RT^2^ Profiler PCR Array for Mouse mTOR Signaling (Qiagen, PAMM-098ZA, Valencia, CA). A complete gene list can be found on the manufacturer’s website. All assays were performed on a ViiA7 Real-Time PCR system (Applied Biosystems, ThermoFisher Scientific, Waltham, MA), in a total volume of 10 μl using 2X SsoFast Evagreen Supermix (BioRad, Hercules, CA) and 50 ng of cDNA per reaction. Standard qPCR settings were used: 95°C for 10 min followed by 40 cycles of 95°C for 15 seconds (sec) then 60°C for 30 sec, followed melt curve analysis. As suggested by the RT^2^ profiler manual, we adjusted the ramp rate to 1°C/sec. All controls including housekeeping genes, positive controls for amplification, and controls for genomic DNA contamination were included as standards in the array.

qPCR data was analyzed using Microsoft Excel. Relative expression was calculated using the ΔΔCt method [39]. Genomic contamination was negligible in all samples. To control for input variations, ΔCt was calculated by comparing the Ct of each gene of interest (GOI) to the average Ct of the 5 housekeeping genes (*Actb*, *B2m*, *Gapdh*, *Gusb*, *Hsp90ab1*) for that sample. ΔΔCt was then calculated relative to expression compared to that seen in the littermate control. Average relative expression (RQ), or fold change (2-ΔΔCt), over controls is shown in Figure S3. All error bars depict RQmax and RQmin, which represent the maximum and minimum limits of possible RQ values based on the standard error of the ΔCt values. The grey line at y = 1 represents the controls.

## QUANTIFICATION AND STATISTICAL ANALYSIS

All data are reported as mean ± standard deviation (S.D.). Statistically significant differences were determined using one-way ANOVA for all experiments with more than two groups but only one dependent variable. Similarly, two-way ANOVA was used for experiments with multiple groups and two dependent variables. All experiments with only two groups and one dependent variable were compared using an unpaired *t*-test with Welch’s correction, which assumes unequal variance. Figure legends specify which test was used for specific experiments. In all cases * = p<0.05; ** = p < 0.01; *** = p<0.001; and **** = p<0.0001; NS = not significant. A “#” symbol was used to highlight data that was trending towards significance (p<0.1). In all cases, asterisks immediately above a bar indicate the significance of that sample relative to the control sample. If any other comparisons, such as *Dhh^Cre(+)^;Fbxw7^fl/+^* (Het) to *Dhh^Cre(+)^;Fbxw7^fl/fl^* (cKO), were significant, this is indicated with a bar spanning above the two samples being compared with the appropriate asterisks. If not indicated otherwise, the comparison was not significant. In most cases, *Dhh^Cre(+)^;Fbxw7^fl/+^* samples were not statistically distinguishable from *Dhh^Cre(+)^;Fbxw7^fl/fl^*.

